# A probabilistic interpretation of PID controllers using active inference

**DOI:** 10.1101/284562

**Authors:** Manuel Baltieri, Christopher L. Buckley

## Abstract

In the past few decades, probabilistic interpretations of brain functions have become widespread in cognitive science and neuroscience. The Bayesian brain hypothesis, predictive coding, the free energy principle and active inference are increasingly popular theories of cognitive functions that claim to unify understandings of life and cognition within general mathematical frameworks derived from information and control theory, statistical physics and machine learning. The connections between information and control theory have been discussed since the 1950’s by scientists like Shannon and Kalman and have recently risen to prominence in modern stochastic optimal control theory. However, the implications of the confluence of these two theoretical frameworks for the biological sciences have been slow to emerge. Here we argue that if the active inference proposal is to be taken as a general process theory for biological systems, we need to consider how existing control theoretical approaches to biological systems relate to it. In this work we will focus on PID (Proportional-Integral-Derivative) controllers, one of the most common types of regulators employed in engineering and more recently used to explain behaviour in biological systems, e.g. chemotaxis in bacteria and amoebae or robust adaptation in biochemical networks. Using active inference, we derive a probabilistic interpretation of PID controllers, showing how they can fit a more general theory of life and cognition under the principle of (variational) free energy minimisation under simple linear generative models.

## 1 Introduction

Probabilistic approaches to the study of living systems and cognition are becoming increasingly popular in the natural sciences. In particular for the brain sciences, theories inspired the Bayesian brain hypothesis such as predictive coding, the free energy principle and active inference have been used to explain brain processes including perception, action and higher order functions [15, 33, 31, 22, 12, 28?, 9].

According to these theories, brains, and biological systems in general, should be thought of as Bayesian inference machines, gathering and representing information from the environment into generative models [22, 12, 28]. Such systems in fact appear to estimate the latent causes of their sensory input in a process consistent with a Bayesian inference scheme. In particular it has been suggested that perceptual process can be accounted for in terms of predictive coding models whereby feedforward prediction errors and feedback predictions are combined under a generative model to infer the hidden causes of sensory data [33]. More recent theories have extended this proposal to account also for motor control and behaviour [26, 19]. On this view, behaviour is cast as a process of acting on the world to make sensory data better fit existing predictions. This latter process usually falls under the name of *active inference.*

Modelling approaches inspired by control theory are nowadays established methodologies for instance in psychology [32, 10], and are increasingly popular in fields such as biology [41, 40, 1]. Typically inspired by classical control theory and dynamical system theory, they emphasise regulation and concepts such as set-point control and negative feedback for the study of different aspects of living systems, an approach first introduced with cybernetics [4, 38]. In particular, methods such as PID (Proportional-Integral-Derivative) control have been widely used as they represent a very simple methodology with properties that guarantee robustness to perturbations and noise [41, 40, 1, 34].

The relationship between information/probability theory and control theory has long been recognised, with the first intuitions (as far as the authors are aware) emerging from work by Shannon [36] and Kalman [30]. A unifying view of these two theoretical frameworks is nowadays proposed for instance in stochastic optimal control [37] and active inference [20, 35]. How these ideas can be used to combine traditional concepts of control more commonly applied in biology with frameworks like active inference is still however unclear. Here, to address this, we develop an information theoretic interpretation of PID controllers, a very popular control strategy that works with little prior knowledge of the process to regulate. Starting from ideas proposed by the free energy principle, we will show that simple linear generative models approximating the true dynamics of the environment implicitly implement PID control as a process of active inference.

## 2 The free energy principle

The free energy principle (FEP) was initially introduced by Karl Friston [22] and later elaborated in a series of papers [18, 19, 24]. The FEP is proposed as a unifying theory for biological sciences with roots in information theory, thermodynamics and statistical mechanics. Work on the FEP has so far covered computational neuroscience [16, 17], and behavioural/cognitive neuroscience studies [26, 23]. Furthermore, connections have been implied with theories of biological self-organisation, information theory (e.g. infomax principle), optimal control, cybernetics and economics (e.g. utility theory) among others [19, 20, 21, 35]. According to the FEP, a living system exists only in a limited set of states over time, e.g. a fish can’t survive for long out of water. Biological creatures can thus be seen as systems that minimise the surprisal (or surprise/self-information) of their sensory observations to maintain their existence, e.g. a fish’ observations should be limited to states in the water. Since this surprisal is not directly accessible by an agent [22], variational free energy is proposed as a proxy that can be minimised in its place, acting as an upper bound for such quantity.

In this study we focus on hypotheses and theories linked to the FEP regarding perception and action in agents, in particular predictive coding and active inference. Predictive coding [33] models of information processing in the brain prescribe a way in which top-down and bottom-up information flows could be combined in the cortex under deep generative models and are, on this view, often described a special case of the FEP [19]. Top-down processes provide predictions about sensory input while bottom-up activity carries prediction errors representing the difference between real and predicted sensations. These errors are then used to train a generative model to produce better predictions. The minimisation of prediction errors achieved by updating the predictions of this model to better represent an agent’s sensations corresponds, on this view, to perception. However, one of the main contributions of the FEP is the extension of predictive coding models to include an account of action, known as *active inference* [25, 26], and thus unify perception and action in a single cohesive mathematical framework where differences between the two processes almost vanish.

### 2.1 Active inference and control

Active inference provides a second way in which prediction errors, or free energy, can be minimised. While perceptual inference suppresses prediction errors only by updating predictions of a generative model of the incoming sensations [18, 26], active inference minimises errors also by directly acting on the environment to change sensory input to better accord with existing predictions. If a generative model encodes information about favourable states for an agent, then this process constitutes a way by which an agent can change its environment to better meet its needs. Thus, under the FEP, these two processes of error suppression allow an agent to both perceive and control the surrounding environment.

Most agent-based models implementing the FEP and active inference assume that agents have a deep understanding of their environment and its dynamics in the form of an accurate and detailed generative model. For instance, in [25, 26] the generative model of the agent explicitly mirrors the *generative process* of the environment, i.e. the dynamics of the world the agent interacts with. In recent work we have argued that this needs not be the case [7], especially if we consider simple living systems with limited resources. We intuitively don’t expect an ant to model the entire environment where it forages, performing complex simulations of the world in its brain (cf. the concept of Umwelt [11]). This idea is however common in other work [25, 26, 23, 28], where cognition and perception are presented as processes of inference to the best explanation, encoding an accurate set of parameters and variables of the environment with agents seen as mainly building sophisticated models of their worlds then used for action and behaviour. One often implicit assumption is that *all* the information available to an agent should be encoded (e.g. an ant modelling the entire environment). This is however not reasonable in dynamic and complex environments: when variables and parameters in the world change too rapidly, accurate online inference and learning are implausible [3]. A possible alternative introduces action-oriented models entailing a more parsimonious approach where only task-relevant information is encoded [13, 14]. On this view, agents only model environmental properties that are necessary for their behaviour and their goals.

In this work we present an example of such parsimonious, action-oriented model described in [13, 14], connecting them to methods from classic control theory. We focus in particular on Proportional-Integral-Derivative (PID) control, both extensively used in industry [6] and more recently emerging as a model of robust feedback mechanisms in biology, implemented for instance by bacteria [41], amoeba [40] and gene networks [1], and in psychology [34]. PID controllers are ubiquitous in engineering mostly due to the fact that one needs only little knowledge of the process to regulate. In active inference terms, we will show that this corresponds to linear generative models that only approximate properties of the world dynamics. Specifically, our model will describe linear dynamics for a single hidden state and a linear mapping from the hidden state to an observed variable, representing knowledge of the world that is potentially far removed from the real complexity behind observations and their hidden causes.

## 3 PID control

Proportional-Integral-Derivative (PID) control is one of the simplest set-point controllers, whereby a desired state (i.e. set-point, reference) represents the final goal of the regulation process, e.g. to maintain a room temperature of 23° C. PID controllers are based on closed-loop strategies with a negative feedback mechanism that tracks the real state of the environment. The difference between such state and the target value (e.g. 23^°^ C temperature) produces a prediction error whose minimisation drives the controller, e.g. if the temperature is too high, it is decreased and if too low, it is raised. In mathematical terms:

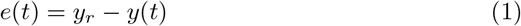

where *e*(*t*) is the error, *y_r_* is the *reference* or set-point (e.g. desired temperature) and *y(t)* is the observed variable (e.g. the actual room temperature).

This mechanism is however unstable in very common conditions, in particular when a steady-state offset is added (e.g. a sudden and unpredictable change in external conditions affecting the room temperature which are not under our control), or when fluctuations need to be repressed (e.g. too many oscillations in the temperature on the trajectory to the desired state may be undesirable). PID controllers deal with both of these problems by augmenting the standard negative feedback architecture, here called *proportional* or *P term*, with an *integral* or *I* and a *derivative* or *D term*, see Fig. 1. The integral term accumulates the prediction error over time in order to cancel out errors due to steady-state input, while minimising the derivative of the prediction error leads to a decrease in the amplitude of fluctuations. The general form of the control signal *u(t)* generated by a PID controller is usually described by:

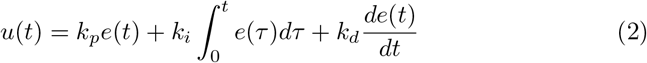

where *e(t)* is again the prediction error and *k_p_,k_i_,k_d_* are the so called proportional, integral and derivative gains respectively, a set of parameters used to tune the relative strength of the P, I and D terms of the controller.

**Fig. 1.**
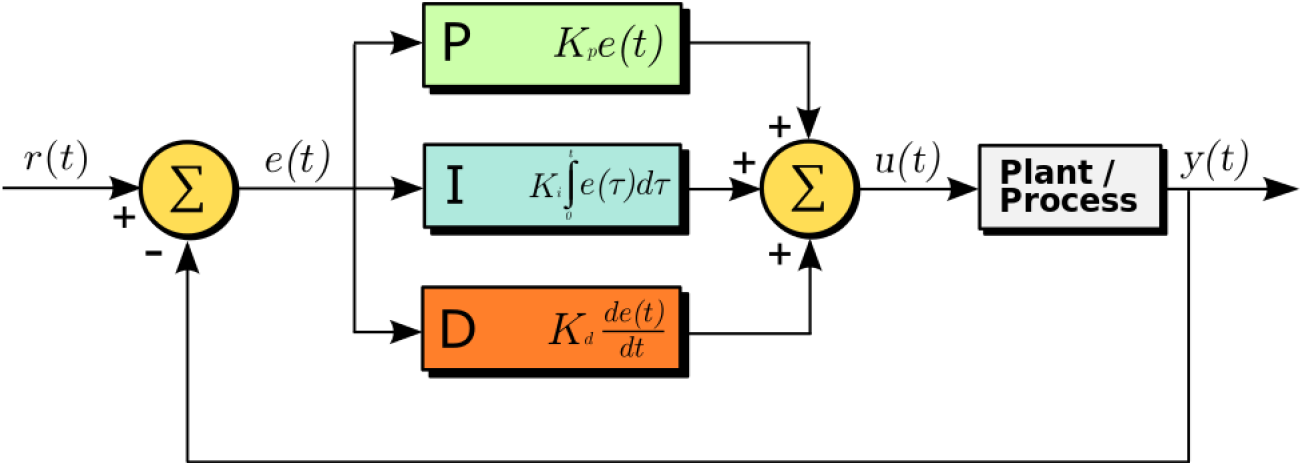
A PID controller [2]. The prediction error *e(t)* is given by the difference between a reference signal *r(t), y_r_* in our formulation, and the output *y(t)* of a process. The different terms, one proportional to the error (P term), one integrating the error over time (I term) and one differentiating it (D term), drive the control signal *u(t).*

The popularity of PID controllers is largely due to: 1) their robustness in the presence of uncertainty, i.e. step disturbances and more in general noise, given by the filtering properties of the I term, and 2) an only approximate model of the dynamics of the process to regulate, based on a linearisation around the target state. This might look incompatible with standard work in active inference formulations suggesting a link to optimal control strategies with perfect models of process/environment, however, we argued previously that this needs not be the case [7]. Indeed one of the main strengths of active inference lies, according to us, in its general formulation and in generative models that do not have to mirror the dynamics of the entire environment.

## 4 PID control as active inference

In this work we will not provide a complete derivation of the active inference scheme, referring to previous treatments [27, 17, 9] for more details. We will begin from the Laplace encoded variational free energy for a univariate case:

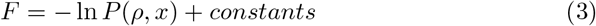

where *ρ* represents the observed sensory input of an agent and *x* encodes the expectation of hidden states in the environment. The remaining constants will not be discussed since they play no role in the minimisation scheme we present.

As previously shown [17, 9], to minimise equation (3) we need to specify the agent’s generative density *P(ρ,x)* = *P(ρ|x)P(x)* introducing a likelihood *P(ρ|x)* and a prior *P(x)* in terms of an agent’s expectations x. These probabilities can be specified by a generative model in the form of a state space model. In particular, to get the integral, proportional and derivative terms of a PID controller, we will use a *generalised* linear state space model of order 2 [17, 9]:

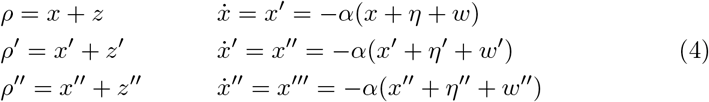

where *ρ* is the observation of the proprioceptive signal and *x* is the estimated hidden state, *α* encodes the decay of *x* (*α* is here assumed to be very large, meaning that the generative model represents a belief in an environment that quickly relaxes to equilibrium), *η* encodes a desired state (e.g. desired temperature) represented mathematically as an exogenous input (or as a prior from higher layers in hierarchical models [17]) and *z,w* are zero-mean Gaussian random variables. As we shall see later, *η* is equivalent to *y_r_* in equation (1). However, unlike standard PID schemes, *η* is here specified as a function of time using generalised coordinates of motion (explained below). Ultimately, this derivation will collapse to a more standard set-point scheme when *η* = *y_r_* and *η*′ = *η*″ = 0. The prime (e.g. *ρ′, ρ′*) indicates the order of generalised coordinate of motion [17, 9], which are introduced to represent non-Markovian continuous stochastic processes [29], in our case *z,w.* One could think of them as quantities conveying information about “velocity” (e.g. *ρ′*), “acceleration” (e.g. *ρ″*), etc. for each variable. Following [17, 9], we then define:

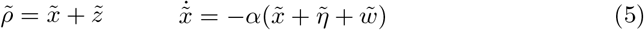

where the tilde sign (e.g. 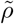) summarises the generalised state, a variable and its higher orders of motion, into a more compact description (e.g. 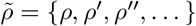).

With the assumption that random variables 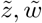 are normally distributed (making the likelihood 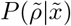 and the prior 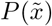 of Gaussian form), the variational free energy reduces to:

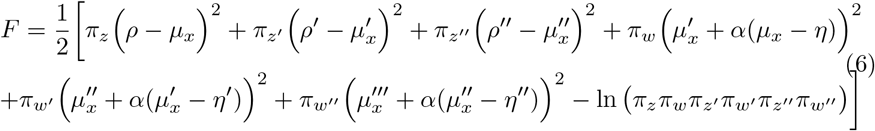

where we used the means 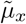 of the estimated hidden states rather than the states 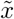 themselves since 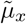 are the only sufficient statistics required for the minimisation of free energy under the assumption of optimal (co)variances of the recognition density (see [17, 9]). 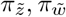 are the precision parameters (inverse variances) of 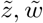 respectively. Following [27, 17], the optimisation of the Laplace encoded free energy can be performed via a standard gradient descent procedure:

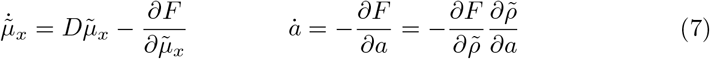

where the two equations prescribe perception and action processes respectively. The first equation includes an extra term 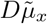 that represents the “mean of the motion” in the minimisation of variables in generalised coordinates of motion [17, 9], with *D* as a differential operator with respect to time, i.e. 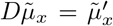. More intuitively, since we are now minimising the components of a generalised state representing a trajectory rather than a static variable, variables are in a moving framework of reference where the minimisation is achieved for 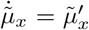 rather than 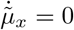. At this point, the mean of the motion becomes the motion of the mean, thereby satisfying Hamilton’s principle of least action [17]. In the second equation, an assumption of active inference is that actions *a* only affect sensory input 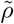 and furthermore that this mapping is known to the agent and enacted as a reflex mechanism, see [26] for discussion. By applying the gradient descent described in equation (7) to our free energy function in equation (6), we then obtain the following update equations for perception:

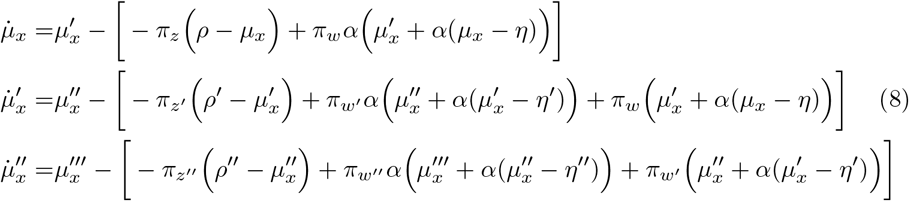

and for action:

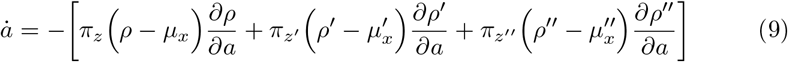

The mapping of these equations to a PID control scheme becomes more clear under a few simplifying assumptions, starting from an agent that will have strong priors (desires) on the causes of its proprioceptive observations. Intuitively, these priors will be used to define actions that change the observations to better fit the agent’s desires. This is implemented in the weighting mechanism of prediction errors by precision parameters in equation (6); see also [26, 8, 7] for similar discussions on precisions and behaviour. Here we want to weight prediction errors on the expected causes, 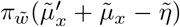, more than the ones on observations, 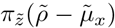. To achieve this, we decrease precisions 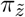 on proprioceptive observations, effectively biasing the gradient descent procedure towards minimising errors on the priors [8]. We then set the decay parameter *α* to a large value, meaning that the agent encodes beliefs in a world that quickly settles to an equilibrium state, with higher orders of generalised motion in each line of equation (8) not considered during the minimisation. Perception is then approximated as:

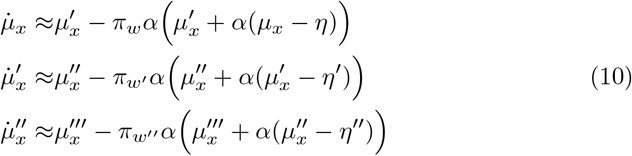

where 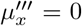 since we truncated our generalised state-space model to order 2 (i.e. anything beyond that is zero-mean Gaussian noise). This system of equations sets, at steady state, the expected hidden states 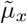 to the priors 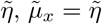.

To minimise free energy in presence of strong priors, the agent will necessarily have to modify its sensory input 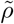 to better match expectations 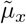 which in turn will be shaped by the priors (i.e. desires) 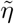. Effectively, the agent “imposes” its desires on the world, driving actions that will minimise the prediction errors arising at the proprioceptive sensory layers. In essence, an active inference agent implements set-point regulation by acting to make its sensations accord with its strong priors/desires. After these assumptions, action can be written as:

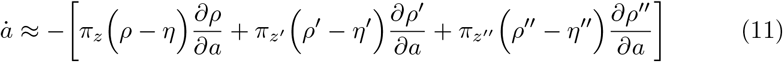

where we assumed 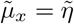, but still need to specify partial derivatives 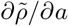. As discussed previously [26], this step also highlights the fundamental differences between the FEP and the more traditional forward/inverse models formulation of control problems [39]. While these derivatives define a form of inverse model, unlike more traditional approaches this does not involve a mapping between actions and hidden states 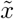 but is cast in terms of sensory data 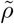 directly. It is claimed that this provides an easier implementation for such an inverse model [20], one that is grounded in an extrinsic frame of reference (i.e. the real world, 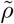) rather than in a intrinsic one in terms of hidden states 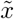. To achieve PID-like control, we finally assume that the agent adopts the simplest (i.e. linear) relationship between its actions (controls) and their effects on sensory input across all generalised coordinates of motion:

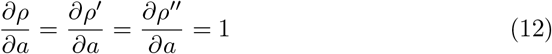

This reflects a very simple reflex-arc-like mechanism that is triggered any time a proprioceptive prediction is made. Intuitively, positive actions increase the values of the sensed variables 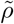, while negative actions decrease them. There is however an apparent inconsistency here that we need to dissolve: the proprioceptive input *ρ* and its higher order states *ρ*′, *ρ*″ are *all* linearly dependent on actions *a* as represented in equation (12). While an action may not change position, velocity and acceleration of a variable in the same way, the goal of an agent is not to perfectly represent its physical reality but just to encode sensorimotor properties that allow it to achieve its goals. In the same way, PID controllers are, in most cases, effective but only approximate solutions for control [5]. This allows us to understand the encoding of an inverse model from the perspective of an agent rather than assuming a perfect, objective mapping from sensations to actions that reflects exactly how actions affect sensory input [26]. This also points at possible investigations of generative/inverse models in simpler living systems where accurate internal models are not needed, and where strategies like PID control are implemented [41, 40, 1]. By combining equation (11) and equation (12), action can then be simplified to:

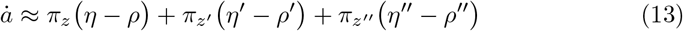

consistent with the “velocity form” or algorithm of a PID controller [5]:

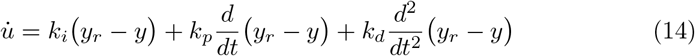

where we removed the explicit dependence on time *t*. Velocity forms are used in control problems where, for instance, integration is provided by an external mechanism outside the controller [5, 6]. This algorithm is often described using discrete systems to avoid the definition of the derivative of random variables, often assumed to be white noise (i.e. Markov processes). In the continuous case, if the variable *y* is a Markov process, its time derivative is in fact not well defined. For this form to exist in continuous systems, *y* must be a smooth process. This effectively drops the Markov assumption of white noise and implements the same definition of analytic (i.e. differentiable) noise and related generalised coordinates of motion we described earlier. The presence of extra prediction errors beyond the traditional negative feedback (proportional term) can in this light be seen as a natural consequence of considering non-Markov processes. To ensure that the active inference implementation approximates the velocity form of PID control we then need to clarify the relationship between generalised coordinates of motion in equation (13) and the differential operators *d/dt*, *d*^2^/*dt*^2^ in equation (14). As pointed out in previous work [27, 9], the two of them are equal at the minimum of the free energy landscape, when the gradient descent has reached its steady state. To simplify our formulation and show this more directly, we could consider the case for *η*′ = *η*″ = 0, defining the more standard set-point control where the desired trajectory collapses to a single point, equivalent in the velocity form to the case where *y_r_* is a constant and *dy_r_/dt* = *d^2^y_r_/dt*^2^ = 0.

## 5 Conclusion

PID controllers are robust controllers used as a model of regulation for noisy and non-stationary processes in different disciplines, from engineering to biology and psychology. They however do not guarantee optimality, so a straightforward interpretation of this control strategy in terms of optimal control is missing. Active inference is often described as an extension of optimal control theory with deep connections to Bayesian inference [20]. While active inference has been proposed as a general mathematical theory of life and cognition according to the minimisation of variational free energy [19], methods such as PID control are still widely adopted as models of biological systems [41, 40, 1]. In this work we proposed a way to connect these two perspectives showing how PID controllers can be seen as a special case of active inference once simplified (i.e. linear) generative models are introduced. The ubiquitous efficacy of PID control may thus reflect the fact that the simplest models of controlled dynamics are first-order approximations to generalised motion. This simplicity is mandated because the minimisation of free energy is equivalent to the maximisation of model evidence, which can be expressed as accuracy minus complexity [19, 14]. On this view, PID control emerges via the implementation of parsimonious (minimum complexity) generative models that are the most effective (maximum accuracy) for a task.

Following our previous work [7], we defined a generative model that only approximates the agent’s environment and showed how under a set of assumptions including analytic (i.e. non-Markovian, differentiable) Gaussian noise and linear dynamics, this model recapitulates PID control. A crucial component of our formulation is the presence of low precision parameters on proprioceptive prediction errors of our free energy function or equivalently, beliefs about high variance of proprioceptive signals. These low precisions play two roles during the minimisation of free energy: (1) they implement control signals as predictions of proprioceptive input influenced by strong priors (i.e. desires) rather than by observations, see equation (11) and [26, 7], and (2) they reflect a belief into the presence of large exogenous fluctuations (low precision = high variance) as part of the observed proprioceptive input. This last point can be seen as the well known property of the Integral term [6] of PID controllers, dealing with unexpected external input (i.e. large exogenous fluctuations). The model represented by derivatives 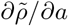 encodes then how actions *a* approximately affect observed proprioceptive sensations 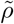, with an agent implementing a sensorimotor mapping that does not match the real dynamics of actions applied to the environment. The generative model we proposed can in general be applied to different tasks, in the same way PID control is used in different problems without specific knowledge of the system to regulate.

In future work we will explore the implications of PID control as active inference for the study of biological systems. In particular we suggest that given our formalisation it is trivial to generalise the set-point definition of PID controllers, based on point attractors, to trajectories (e.g. a reference temperature changing during the day with pre-specified properties such as the rate of change, etc.) using generalised coordinates of motion [27, 9]. We may also be able to provide a Bayes-optimal algorithm for the optimisation of the gains of a PID controller, *k_i_, k_p_, k_d_* (i.e. precision parameters in our free energy formulation, *π_w_, π_w′_, π_w″_*), for which only heuristic methods exist at the moment [5].

## 6 Acknowledgements

The authors would like to thank Karl Friston for thought-provoking discussions and insightful feedback on the final version of this manuscript, and Martijn Wisse and Sherin Grimbergen for important comments on the mathematical derivation.

